# Blockade of SARS-CoV-2 infection in vitro by highly potent PI3K-α/mTOR/BRD4 inhibitor

**DOI:** 10.1101/2021.03.02.433604

**Authors:** Arpan Acharya, Kabita Pandey, Michellie Thurman, Kishore B. Challagundala, Kendra R. Vann, Tatiana G. Kutateladze, Guillermo A Morales, Donald L. Durden, Siddappa N. Byrareddy

## Abstract

Pathogenic viruses like SARS-CoV-2 and HIV hijack the host molecular machinery to establish infection and survival in infected cells. This has led the scientific community to explore the molecular mechanisms by which SARS-CoV-2 infects host cells, establishes productive infection, and causes life-threatening pathophysiology. Very few targeted therapeutics for COVID-19 currently exist, such as remdesivir. Recently, a proteomic approach explored the interactions of 26 of 29 SARS-CoV-2 proteins with cellular targets in human cells and identified 67 interactions as potential targets for drug development. Two of the critical targets, the bromodomain and extra-terminal domain proteins (BETs): BRD2/BRD4 and mTOR, are inhibited by the dual inhibitory small molecule SF2523 at nanomolar potency. SF2523 is the only known mTOR PI3K-α/(BRD2/BRD4) inhibitor with potential to block two orthogonal pathways necessary for SARS-CoV-2 pathogenesis in human cells. Our results demonstrate that SF2523 effectively blocks SARS-CoV-2 replication in lung bronchial epithelial cells *in vitro*, showing an IC_50_ value of 1.5 µM, comparable to IC_50_ value of remdesivir (1.1 µM). Further, we demonstrated that the combination of doses of SF2523 and remdesivir is highly synergistic: it allows for the reduction of doses of SF2523 and remdesivir by 25-fold and 4-fold, respectively, to achieve the same potency observed for a single inhibitor. Because SF2523 inhibits two SARS-CoV-2 driven pathogenesis mechanisms involving BRD2/BRD4 and mTOR signaling, our data suggest that SF2523 alone or in combination with remdesivir could be a novel and efficient therapeutic strategy to block SARS-CoV-2 infection and hence be beneficial in preventing severe COVID-19 disease evolution.

**One Sentence Summary:** Evidence of *in silico* designed chemotype (SF2523) targeting PI3K-α/mTOR/BRD4 inhibits SARS-CoV-2 infection and is highly synergistic with remdesivir.

## INTRODUCTION

Coronavirus disease 2019 (COVID-19) is caused by a novel coronavirus named severe acute respiratory syndrome coronavirus 2 (SARS-CoV-2) (*1*). This is a novel zoonotic single-stranded RNA virus that belongs to the betacoronavirus family, which also includes severe acute respiratory syndrome coronavirus (SARS-CoV) and middle east respiratory syndrome coronavirus (MERS-CoV) (*2*). The ongoing global pandemic of SARS-CoV-2 was first reported from Wuhan, China, in January 2020. As of March 01, 2021, globally, more than 114.30 million confirmed cases and 2.53 million deaths had been reported due to COVID-19 (https://coronavirus.jhu.edu/map.html).

SARS-CoV-2 utilizes host cell angiotensin-converting enzyme 2 (ACE2) as the primary receptor for entry along with several other accessory factors that include TMPRSS2, neuropilin-1, PIKFyve kinase, and CD147, among others (*3-8*). After internalization, cathepsin L cleaves the S protein and aids in releasing the viral genome in the host cell cytoplasm (*3*). Inside the cell, virus utilizes the host cell machinery to replicate its positive-sense RNA genome and process the polyproteins to generate functional structural and nonstructural proteins. Then, the viral genome and structural proteins are assembled into progeny viruses, travel to the plasma membrane and are released utilizing lysosomal exocytosis pathways (*9*). During this replication cycle, several viral proteins like NSP1, NSP3, and ORF7a/b suppress the type 1 interferon (IFN) mediated host immune response (*10*), while others activate Toll-like receptors (TLRs) that stimulate the release of proinflammatory cytokines like IL-1β, IL-2, IL-6, IL-7, GCSF, IFN-γ, and TNF-α (*11*). The pathophysiology of COVID-19 follows a biphasic pattern. In contrast, the initial acute phase of infection is manifested by viral infection-driven symptoms including sore throat, fever, fatigue, dry cough, and/or diarrhea (*12*). In severely ill patients, this is followed by an immunopathologic phase that includes the development of acute respiratory distress syndrome (ARDS), systemic inflammation, and cytokine storm, which is responsible for multiple organ failure and a higher rate of fatalities (*1*). Patients with certain comorbid conditions like hypertension, cardiovascular disease, diabetes, and chronic obstructive pulmonary disorders (COPD) also have a higher probability of developing acute severe respiratory distress syndrome (ARDS). Moreover, a subpopulation of patient’s develops neurological disease manifestations like loss of taste and smell, dizziness, confusion, ataxia, seizures and Guillain-Barré Syndrome (GBS) have also been reported (*13*). Among the patients who develop ARDS, many of them succumbed to the disease due to the development of pulmonary fibrosis (*14*). These data suggest that the addition of a currently available or upcoming anti-fibrotic agent for the management of critically ill COVID-19 patients would be beneficial to reduce the rate of fatalities and long-term consequences of pulmonary fibrosis.

Although some vaccine candidates received emergency use authorization (EUA) from U.S. Food and Drug Administration (USFDA) to control the rapid spread of the virus, it is anticipated that it will take several months to years to vaccinate the majority of the population (*15*). Again the long-term safety, efficacy, and durability of these vaccines remain unknown specifically for the ‘special’ populations that include children, pregnant women, individuals with comorbid conditions that may impact immune response against a vaccine (*16*). Along with that, preliminary studies indicate that the emerging new strains of SARS-CoV-2 from South Africa having Lys417Asn, Glu484Lys, and Asn501Tyr substitutions in the receptor-binding domain, and Leu18Phe, Asp80Ala, Asp215Gly, 242–244del, and Arg246Ile in the N-terminal domain of spike protein have reduced susceptibility to neutralization by convalescent plasma therapy (*17*). These results question the efficacy of the spike protein-based vaccines against new strains of the virus and further support the fact that this virus will eventually enter an endemic stage with world-wide distribution and high-level mortality. Therefore, other strategies to reduce SARS-CoV-2 and related mutant strain associated morbidity and mortality are urgently needed. These factors underscore the urgent development of an effective therapeutic regimen that will control the rapid spread of the virus and reduce the high rate of fatalities among severely ill patients. From the outset of this pandemic, multiple therapeutic strategies have been tested that are at different stages of development and clinical trials with mixed responses (*18*). The majority of which includes SARS-CoV-2 fusion/entry inhibitors (*19*), RNA-dependent RNA polymerases (RdRps) inhibitors (*20*), viral protease inhibitors (*21*), several classes of immune-suppressive agents to control systemic hyperactive inflammatory response (*22*), and cytokine storm and convalescent plasma treatment (*23*).

At present, remdesivir (GS-5734) is the only US FDA-approved antiviral agent in use to treat COVID-19 (*24*). It is a prodrug of a monophosphate nucleotide analog that selectively inhibits RNA-dependent RNA polymerase of SARS-CoV-2 (*25*). A randomized, double-blind, placebo-controlled clinical trial reported shorter hospital stay for patients with COVID-19 when treated with remdesivir than placebo controls (ClinicalTrials.gov: NCT04280705) (*26*). However, another multicenter, randomized clinical trial conducted by WHO did not find any superior effect of remdesivir on hospitalized COVID-19 patients (ClinicalTrials.gov: NCT04315948) (*27*). Therefore, in the absence of a fully effective drug in the prevention of COVID-19, it is warranted to identify and validate more compounds that, individually or in combination with other molecules, will have a profound antiviral efficacy against COVID-19 (*28*).

Recently, Gordon et al. (*29*) used a proteomic approach to explore the interactions of 26 out of 29 SARS-CoV-2 proteins with cellular targets in human cells and identified 67 potential interactions and targets for drug development. By molecular pathway analysis, they demonstrated an interaction of BRD2/BRD4 with E protein of SARS-CoV-2. In another study Tian et al. using CRISPRi screening shown that BRD2 inhibition down regulate the ACE2 expression which block the entry of SARS-CoV-2 within host cells as well as control hyperactive immune response in COVID-19 patients through down regulations of ISGs (*30*). Therefore, they proposed BRD2 as a potential target for development of therapeutics against SARS-CoV-2. Two of the critical host factors responsible for pathogenesis of SARS-CoV-2, the bromodomain and extra-terminal domain proteins (BETs): BRD2/BRD4 and mTOR, are inhibited by the dual inhibitory small molecule SF2523 developed by our group previously. In the present study our *in vitro* data using a human bronchial epithelial cell line (UNCN1T) shows that SF2523 effectively blocks SARS-CoV-2 replication with IC_50_ values comparable to that of remdesivir. We also found that the combination of doses of SF2523 and remdesivir is highly synergistic. There is a 25-fold dose reduction for SF2523 and a 4-fold dose reduction for remdesivir, respectively, when the two compounds are used in combination and they maintain similar potency as a single compound. Future studies in suitable animal models are needed to establish the clinical efficacy of SF2523 alone or a fixed-dose combination of SF2523 and remdesivir as a novel treatment of COVID-19.

## RESULTS

### Molecular interaction of SF2523 with PI3K-α and BRD4

In a recent study using proteomic analysis, Gordon et al. (*29*) identified 67 host factors that interact with SARS-CoV-2 proteins and represent potential drug development targets. Two of the potential host targets identified, BETs BRD2/BRD4 and mTOR, are inhibited by SF2523. Molecular modeling and structural analysis of predicted molecular interactions of dual BET (BRD2/4)/PI3K-mTOR inhibitor SF2523 with the catalytic domain of PI3K-α are displayed in the top panel of **Fig. 1**. The molecular interactions of SF2523 binding to BRD4-BD1 binding pocket were observed from a 1.8 angstrom resolution crystal structure (PDB ID: 5U28) displayed in the bottom panel of **Fig. 1**. In a cell-free system, the binding potential of SF2523 with its potential targets is shown below the bottom panel of **Fig 1**.

**Fig. 1.**
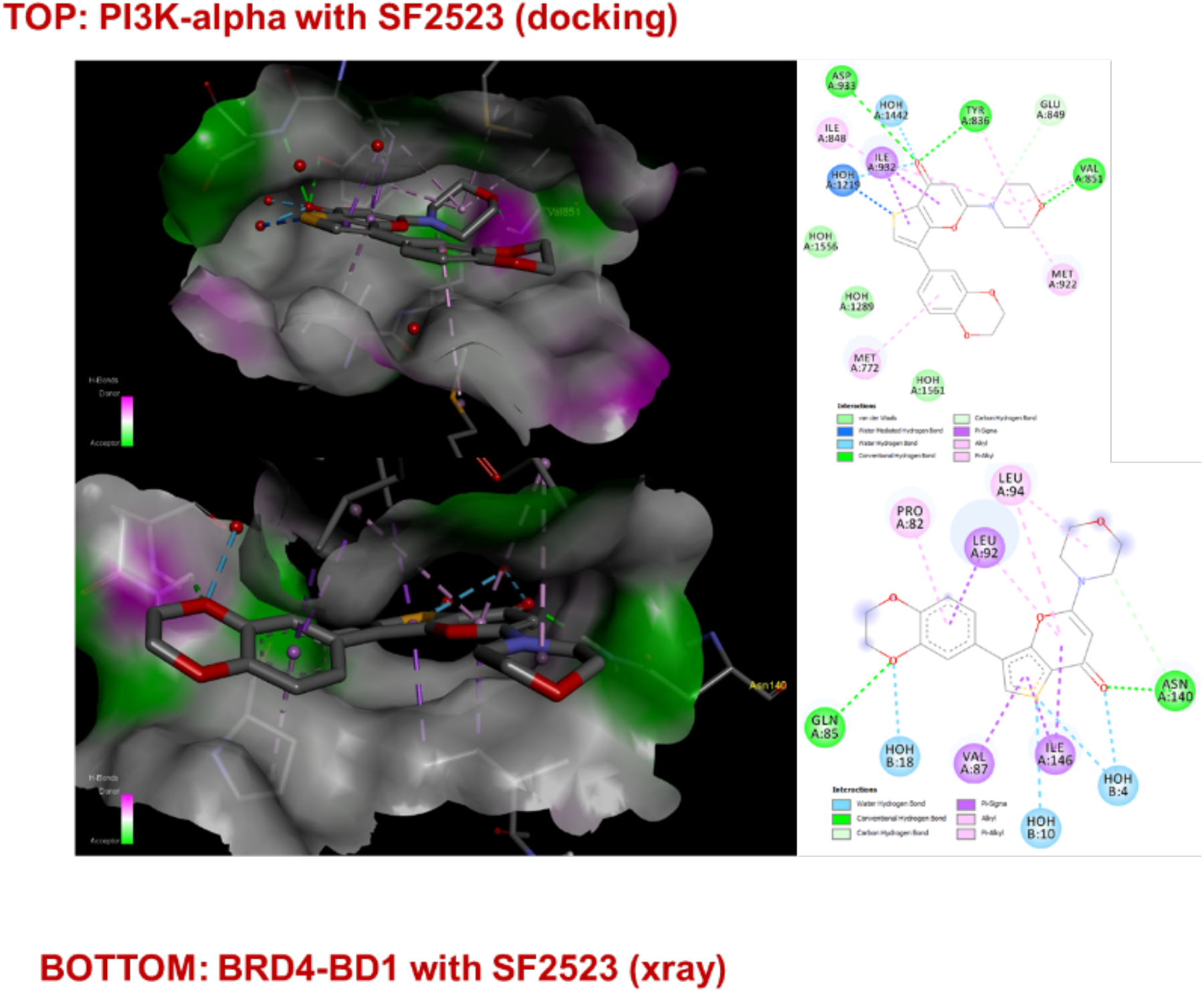
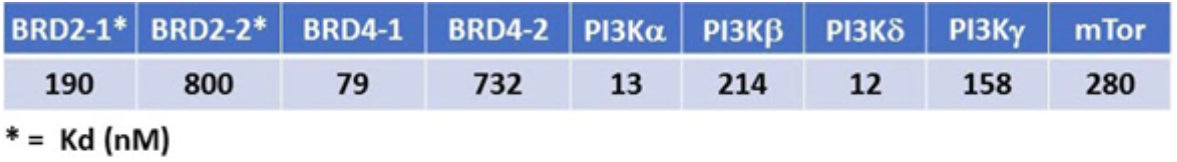
Predicted binding (docking) of SF2523 in PI3K-α catalytic domain (upper panel) and observed binding in crystal structure of SF2523 bound to BRD4-BD1 acetyl lysine binding site (1.8 Angstroms resolution) (bottom panel). *In vitro* (enzymatic) potency of SF2523 against targets.

### Antiviral efficacy of SF2523 and remdesivir

This study evaluates SF2523 and remdesivir (GS-5734) antiviral efficacy against SARS-CoV-2 in Vero-STAT1 knockout cells and UNCN1T cells. STAT1 is a transcription factor essential for interferon-mediated host-cell antiviral response. The STAT1 knockout cells are highly susceptible to viral infection due to the absence of cellular antiviral response and used as control for robust viral infection in this study. On the other hand, UNCN1T is a human bronchial epithelial cell line and expected to demonstrate the *in vivo* pathogenesis of SARS-CoV-2 in the lung and serves as a good *in vitro* model for testing efficacy of antiviral compounds against SARS-CoV-2.

Both cell types were treated with SF2523 or remdesivir and their IC_50_ values were determined 24 hrs, and 48 hrs post-infection. The replication kinetics were determined from the culture supernatant of SARS-CoV-2 infected UNCN1T and Vero-STAT1 knockout cells in the presence of increasing concentrations of SF2523 and remdesivir and are described in **Fig. 2A** and **Fig. 2B**, respectively. Both compounds show a substantial reduction in SARS-CoV-2 viral load 24 hrs post-infection. Based on SARS-CoV-2 viral loads 24 hrs post infection, SF2523 showed potential antiviral activity and inhibited infection in UNCN1T cells with an IC_50_ of 1.52 μM and remdesivir with an IC_50_ of 1.06 μM. In Vero-STAT1 knockout cells, SF2523 has an IC_50_ of 1.02 μM and remdesivir has an IC_50_ of 1.03 μM (**Fig. 2C and 2D**). SARS-CoV-2 replication kinetics 48 hrs post-infection in SF2523 and remdesivir treated UNCN1T and Vero-STAT1 knockout cells are described in **Fig. S1A** and **Fig. S1B**, respectively. SF2523 and remdesivir have an IC_50_ value of 1.58 μM and 2.75 μM in UNCN1T cells (**Fig. S1C)** and IC_50_ of 3.22 μM and 0.76 μM in Vero-STAT1 knockout cells (**Fig. S1D**), respectively.

**Fig 2.**
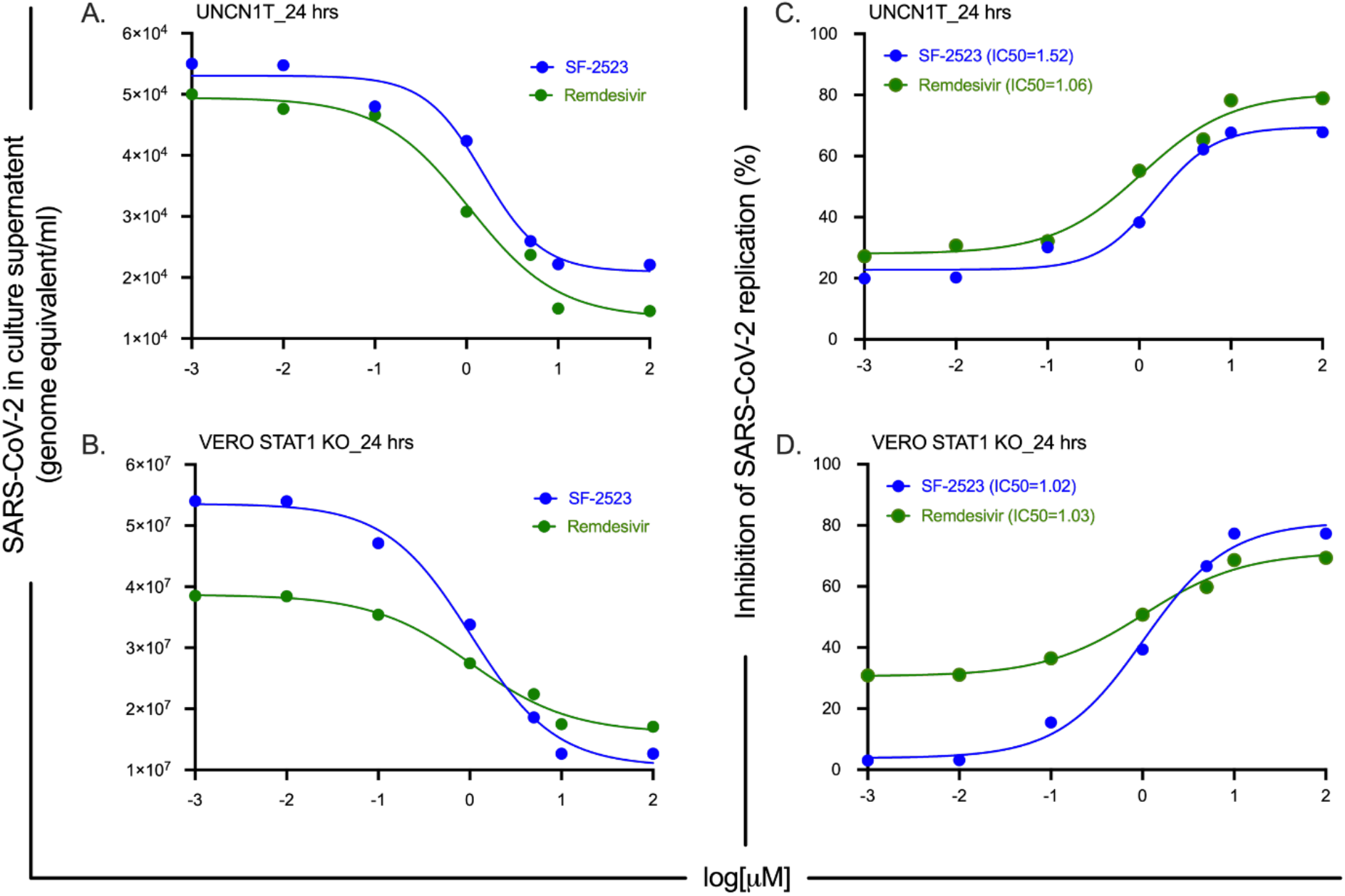
SARS-CoV-2 viral loads and dose-response curve in SF2523 and remdesivir treated and SARS-CoV-2 infected UNCN1T cells and Vero-STAT1 knockout cells. (A & B) Real-time quantitative PCR analysis of SARS-CoV-2 genome equivalent per ml of culture supernatant after 24 hrs post-infection in SF2523 (in blue) and remdesivir (in green) treated UNCN1T cells and Vero-STAT1 knockout cells with indicated drug concentrations, respectively. (C & D) SF2523 (in blue) and remdesivir (in green) dose response curve by percentage inhibition of SARS-CoV-2 replication 24 hrs post infection in UNCN1T cells and Vero-STAT1 knockout cells with indicated drug concentrations, respectively.

### Dose-response relationship and synergy of SF2523 with remdesivir

After testing SF2523 and remdesivir separately, we evaluated a series of combinations of doses of SF2523 and remdesivir with concentrations above and below the IC_50_ determined above. In UNCN1T cells at 24 hrs post-infection, we observed the best inhibitory dose combination. An IC_50_ value of 0.62 μM was observed for remdesivir when combined with 0.1 µM SF2523 (**Fig. 3A**). Vice versa, we saw the inhibition of viral replication at a fixed remdesivir dose of 0.05 μM and an IC_50_ value of 1.21 μM for SF2523. (**Fig. 3B**). The use of multiple drugs with a different modes of action and synergistic effects is a standard for treatment of infectious diseases to increase the overall efficacy of the therapy and reduce dosage of individual drugs while also reducing the toxicity level and the chances of development of drug resistance (*31*). Based on mass action law and the classic model of enzyme-substrate inhibitor interactions, the median-effect equation was proposed by Chou (*32*), which describes the dose-effect relationship as (Fa/Fu) = (D/Dm)^m^ or log(Fa/Fu) = mlog(D) – mlog(Dm). Where the slope (m) of the lines signifies the shape (m = 1, >1, and <1 signify hyperbolic, sigmoidal, and flat sigmoidal dose-effect curves, respectively). Fa is defined as percentage inhibition of viral growth at dose D, and Fu is the fraction that remains unaffected (i.e., Fu = 1 - Fa). The antilog of x-intercept, where Fa/Fu = 1 or log(Fa/Fu) = 0, gives the Dm value. [log(Dm)] signifies the potency of the drugs (where Dm stands for median effective dose or IC_50_ concentration: The concentration required to inhibit 50% growth of the virus). By extrapolating the median-effect equation for multiple drugs, Chou and Talalay proposed the combination index (CI) theorem to quantify synergism and antagonism between drug combinations (*33*). Low CI values with increased Fa values indicate superior compatibility and high synergism between drugs combination (based on Chou and Talalay Combination Index Theorem, CI < 1, = 1, and >1 show synergism, additive effect, and antagonism, respectively). A plot of CI values at different fraction affected (Fa) levels referred to as Fa-CI plot or the Chou-Talalay plot.

**Fig 3.**
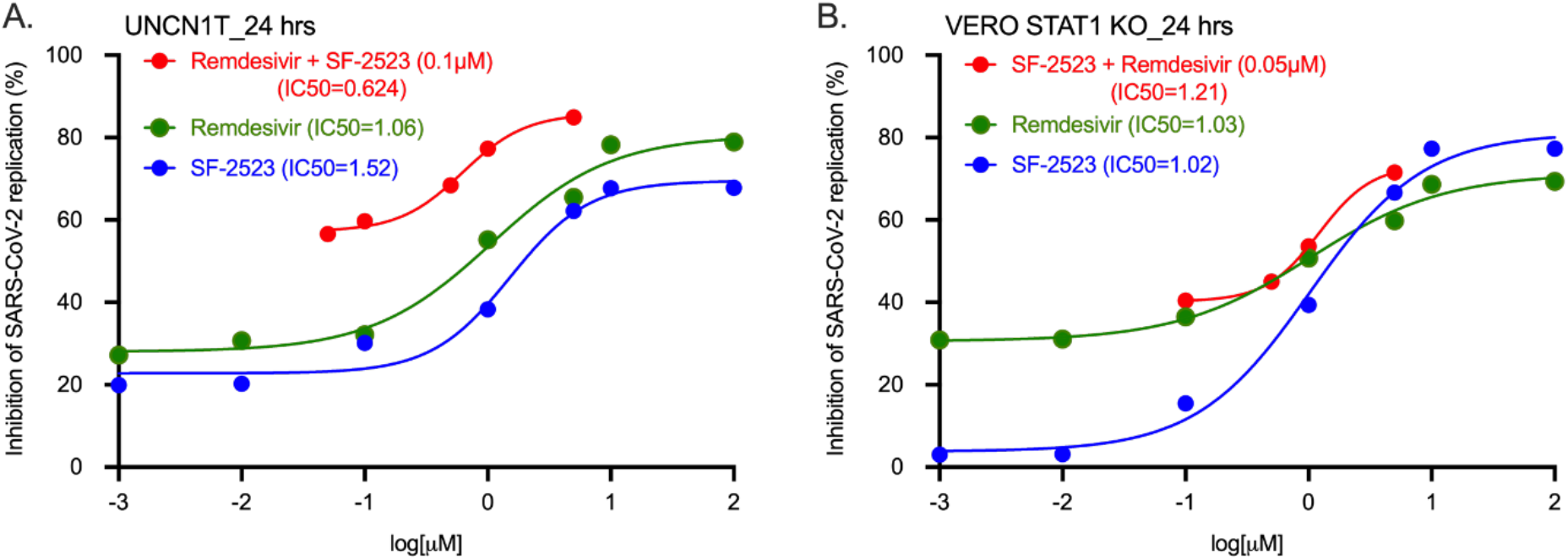
Dose-response curve for single and combination of SF2523 and remdesivir in SARS-CoV-2 infected UNCN1T cells and Vero-STAT1 knockout cells 24 hrs post infection. (**A**) Dose response curve of remdesivir (green: IC_50_ = 1.06 μM), (blue: IC_50_ = 1.52 μM), and remdesivir with a fixed dose combination of SF2523 (0.1 μM) (red: IC_50_ = 0.625) by percentage inhibition of SARS-CoV-2 replication 24 hrs post infection in UNCN1T cells; (**B**) Dose response curve of remdesivir (green: IC_50_ = 1.03μM), (blue: IC_50_ = 1.02 μM), and SF2523 with a fixed dose combination of remdesivir (0.05μM) (red: IC_50_ = 1.21) by percentage inhibition of SARS-CoV-2 replication 24 hrs post infection (IC_50_ = 1.02 μM) in Vero-STAT1 knockout cells.

Using CompuSyn, we generated the Median-Effect plot (Chou plot) of remdesivir (red line), SF2523 (blue line), and the fixed-dose combination of remdesivir and SF2523 (green line) for SARS-CoV-2 infected UNCN1T cells 24 hrs post infection as described above (**Fig. 4A**). Analysis of the Median-Effect, where the X-intercept of the lines indicates potency, shows that the combination dose is more potent than the individual doses of the compounds. In contrast, the Chou-Talalay plot (**Fig. 4B**) indicates the synergistic effect of SF2523 and remdesivir. The dose-response percent inhibition matrix of single and combined treatment of remdesivir and SF2523 was computed using SynergyFinder v.2 (**Fig. 4C**). The 3-D interaction landscape between remdesivir and SF2523 was calculated based on Loewe additive model using SynergyFinder v.2 in SARS-CoV-2 infected UNCN1T cells 24 hrs post-infection (Loewe synergy score 34.88; with most synergistic area score of 40.24) (**Fig 4D**). Synergy maps highlight synergistic and antagonistic dose regions in red and green colors, respectively. Although there is no defined threshold, drug combinations with a synergy scores above -10 is considered likely antagonistic, and a score between -10 to 10 is deemed a possible additive. Any score above 10 is considered synergistic.

**Fig 4.**
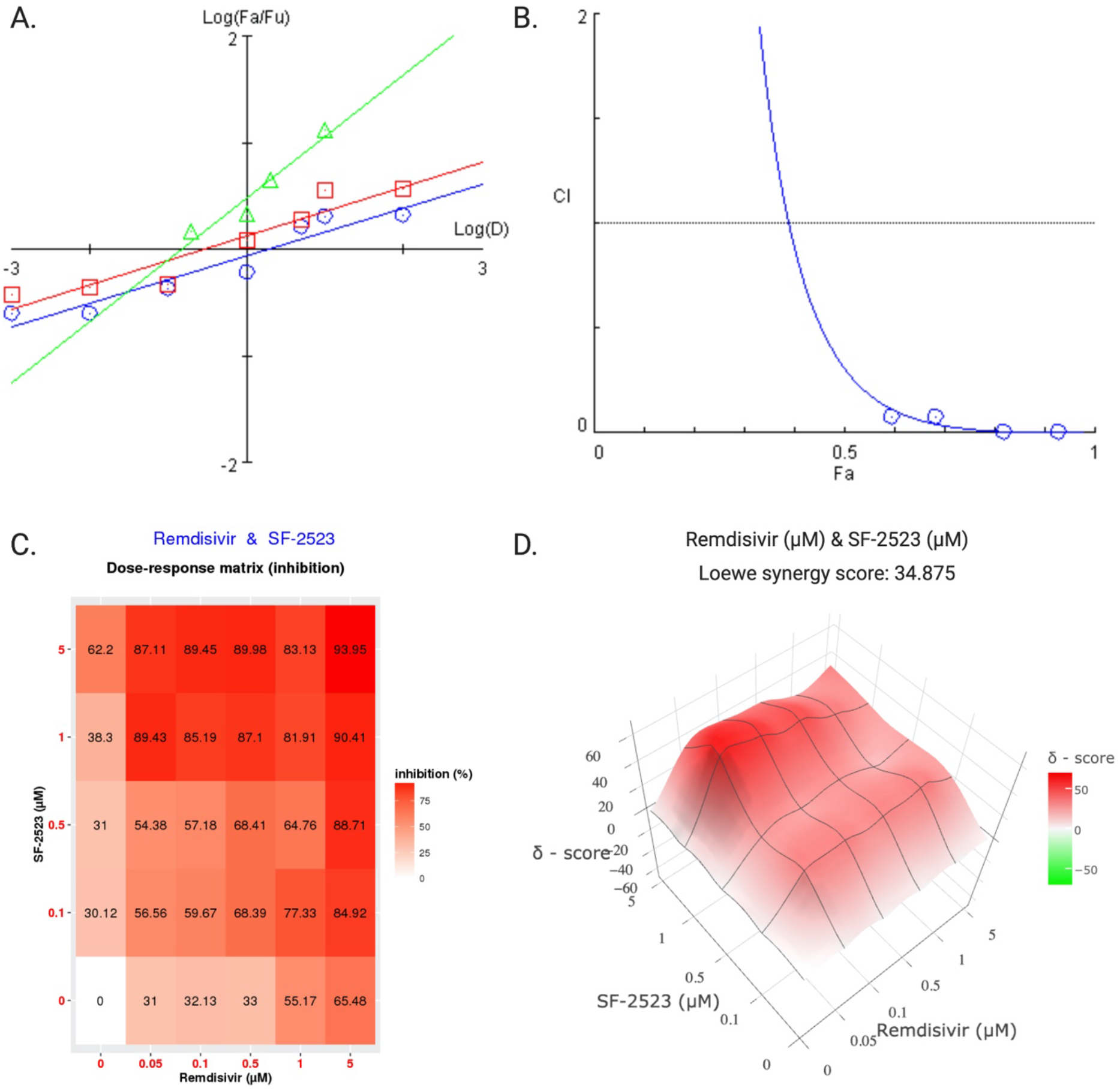
Synergistic effect of remdesivir and SF2523 combined treatment against SARS-CoV-2 infected UNCN1T cells 24 hrs post infection. **(A)** The Median-Effect plot (Chou plot) of remdesivir (the red line with square data points, SF2523 (blue line with circular data points) and fixed dose combination of remdesivir and SF2523 (green line with triangular data points). The median-Effect equation, that describes the dose-effect relationship is given by (Fa/Fu) = (D/Dm)^m^ or log(Fa/Fu) = m log(D) – m log(Dm). The slope (m) of the lines signifies the shape (m = 1, >1, and <1 signify hyperbolic, sigmoidal, and flat sigmoidal dose-effect curves respectively). Fa is defined as percentage inhibition of viral growth at dose D and Fu is the fraction that remain unaffected (i.e., Fu = 1 - Fa). The antilog of x-intercept, where Fa/Fu = 1 or log(Fa/Fu) = 0, gives the Dm value. [log(Dm)] signifies the potency of the drugs (where Dm stands for median effective dose or IC_50_ concentration: The concentration required to inhibit 50% growth of the virus). (**B**) Combination index plot or Chou-Talaly plot or Fa-CI plot: A plot of CI (combination index) on the Y-axis vs Fa on X-axis for fixed dose combination of remdesivir and SF2523 in SARS-CoV-2 infected UNCN1T cells (blue line with circular data points); Low CI values with increase in Fa values indicates superior compatibility and high synergism between drugs combination (based on Chou and Talalay Combination Index Theorem, CI < 1, = 1, and >1 indicate synergism, additive effect, and antagonism, respectively). (**C**) Dose-response percent inhibition matrix of single and combined treatment of remdesivir and SF2523 in SARS-CoV-2 infected UNCN1T cells 24 hrs post infection. (**D**) 3-D interaction landscape between remdesivir and SF2523 calculated based on Loewe additive model using SynergyFinder v.2 in SARS-CoV-2 infected UNCN1T cells 24 hrs post infection (Loewe synergy score 34.88; with most synergistic area score of 40.24). Synergy maps highlight synergistic and antagonistic dose regions in red and green colors, respectively. Although there is no defined threshold, for drug combinations with a synergy scores above -10 is considered likely antagonistic, score between - 10 to 10 is considered likely additive and score above 10 is considered synergistic.

Similarly, for SARS-CoV-2 infected Vero-STAT1 knockout cells treated with different combination doses of remdesivir and SF2523, the Median-Effect plot (Chou plot) of remdesivir (red line), SF2523 (blue line), and the fixed-dose combination of remdesivir and SF2523 (green line) is described in **Fig. 5A**, the Chou-Talalay plot (**Fig. 5B**) indicates a synergistic effect of SF2523 and remdesivir. The dose-response percent inhibition matrix of single and combined treatment of remdesivir and SF2523 was computed using SynergyFinder v.2 (**Fig. 5C**) and the 3-D interaction landscape between remdesivir and SF2523 was calculated based on a Loewe additive model using SynergyFinder v.2 in SARS-CoV-2 infected UNCN1T cells 24 hrs post-infection (Loewe synergy score -0.20; with most synergistic area score of 7.67) (**Fig. 5D**). **Table 1** indicates the median effective dose of SF2523, remdesivir, and their fixed-dose combinations. Whereas **Table 2** shows the CI values at ED_50_, ED_75_, ED_90_, and ED_95_ in VERO-STAT1 knockout and UNCN1T cells, which suggests a moderate to strong synergistic effect of the drug combinations. In Vero-STAT1 KO cells at Fa 0.5, we obtained a dose reduction index (DRI) of 11.56 for SF2523 and 2.35 for remdesivir, respectively (with Fa for combination 0.452; 2.617 for SF2523 and 0.533 for remdesivir). On the other hand, in UNCN1T cells, we obtained a dose reduction index (DRI) of 25.33 for SF2523 and 3.75 for remdesivir, respectively, (with Fa for combination 0.145; 1.836 for SF2523 and 0.272 for remdesivir).

**Fig 5.**
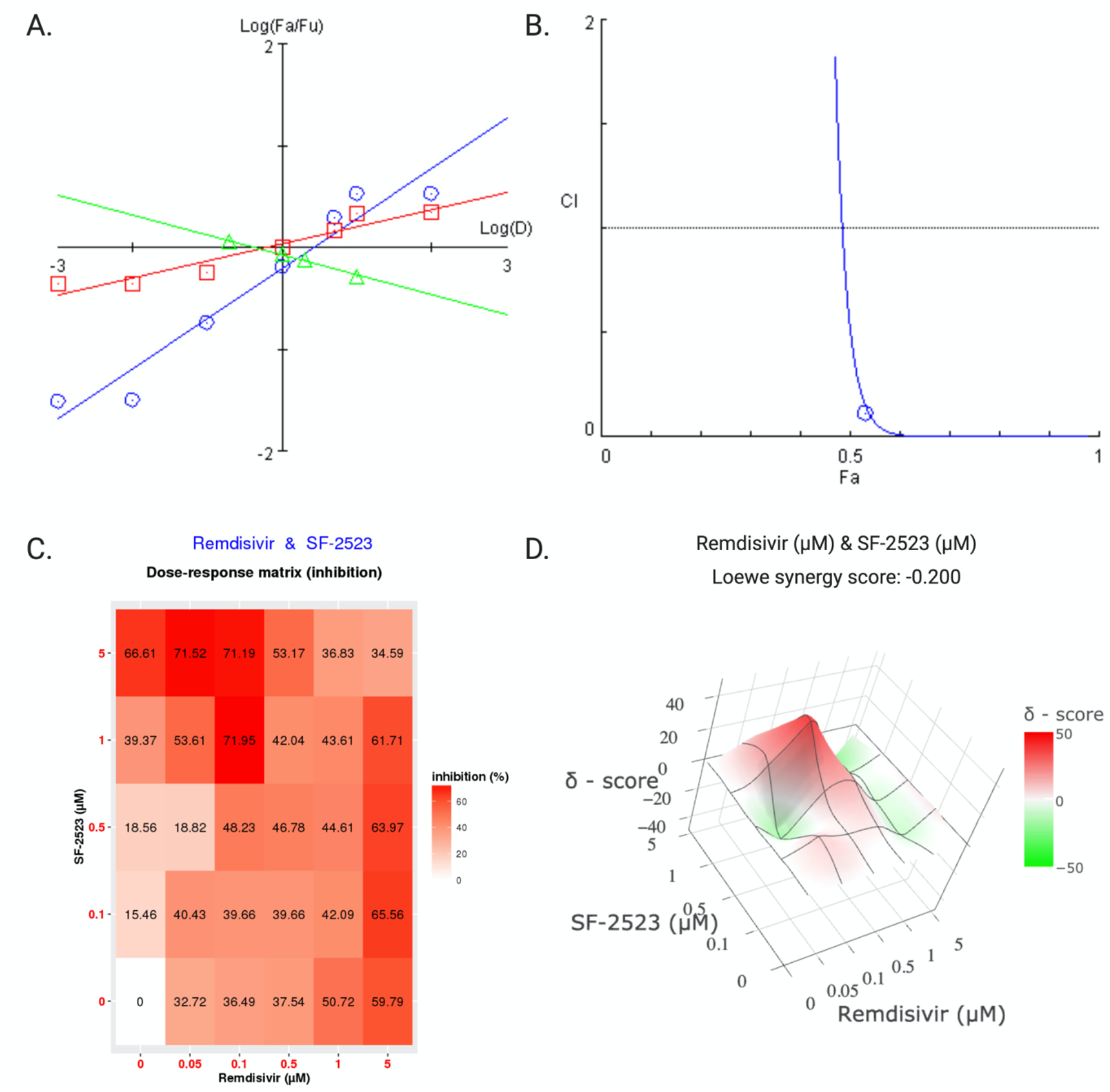
Synergistic effect of remdesivir and SF2523 combined treatment against SARS-CoV-2 in Vero-STAT1 knockout cells 24 hrs post infection. **(A)** The Median-Effect plot (Chou plot) of remdesivir (the red line with square data points, SF2523 (blue line with circular data points) and fixed dose combination of remdesivir and SF2523 (green line with triangular data points). The median-Effect equation, that describes the dose-effect relationship is given by (Fa/Fu) = (D/Dm)^m^ or log(Fa/Fu) = m log(D) – m log(Dm). The slope (m) of the lines signifies the shape (m = 1, >1, and <1 signify hyperbolic, sigmoidal, and flat sigmoidal dose-effect curves respectively). Fa is defined as percentage inhibition of viral growth at dose D and Fu is the fraction that remain unaffected (i.e., Fu = 1 - Fa). The antilog of x-intercept, where Fa/Fu = 1 or log(Fa/Fu) = 0, gives the Dm value. [log(Dm)] signifies the potency of the drugs (where Dm stands for median effective dose or IC_50_ concentration: The concentration required to inhibit 50% growth of the virus). (**B**) Combination index plot or Chou-Talaly plot or Fa-CI plot: A plot of CI (combination index) on the Y-axis vs Fa on X-axis for fixed dose combination of remdesivir and SF2523 in SARS-CoV-2 infected Vero-STAT1 knockout cells (blue line with circular data points); Low CI values with increase in Fa values indicates superior compatibility and high synergism between drugs combination (based on Chou and Talalay Combination Index Theorem, CI < 1, = 1, and >1 indicate synergism, additive effect, and antagonism, respectively). (**C**) Dose-response percent inhibition matrix of single and combined treatment of remdesivir and SF2523 in SARS-CoV-2 infected Vero-STAT1 knockout cells 24 hrs post infection. (**D**) 3-D interaction landscape between remdesivir and SF2523 calculated based on Loewe additive model using SynergyFinder v.2 in SARS-CoV-2 infected Vero-STAT1 knockout cells 24 hrs post infection (Loewe synergy score -0.20; with most synergistic area score of 7.67). Synergy maps highlight synergistic and antagonistic dose regions in red and green colors, respectively. Although there is no defined threshold, for drug combinations with a synergy scores above -10 is considered likely antagonistic, score between -10 to 10 is considered likely additive and score above 10 is considered synergistic.

**Table 1:** The median effective dose of individual compounds and their combination

| **Vero-STAT1 knockout Cells** |  |  |  |
| --- | --- | --- | --- |
| **Drug/Combo** | **Dm** | **m** | **r** |
| **SF2523 (S)** | 2.617 | 0.492 | 0.971 |
| **Remdesivir (R)** | 0.533 | 0.167 | 0.955 |
| **Combination (SRF)** | 0.452 | -0.197 | -0.994 |
| **UNCN1T Cells** |  |  |  |
| **Drug/Combo** | **Dm** | **m** | **r** |
| **SF2523 (S)** | 1.836 | 0.222 | 0.953 |
| **Remdesivir (R)** | 0.272 | 0.231 | 0.949 |
| **Combination (SRF)** | 0.145 | 0.577 | 0.967 |

Dm: The median effective dose; m: The slope of the median effect (ME) plot; r: The linear correlation coefficient of ME plot.

**Table 2:** The CI values at different simulated effective dose of fixed dose combination of SF2523 and remdesivir

| **CI values** |  |  |  |  |
| --- | --- | --- | --- | --- |
|  | **ED50** | **ED75** | **ED90** | **ED95** |
| Vero-STAT1 knockout cells | 0.5105 | 3.75E-05 | 1.44E-08 | 7.10E-11 |
| UNCN1T cells | 0.3054 | 0.0173 | 9.91E-04 | 1.42E-04 |

### NMR studies involving SF2523 and BRD4

We note that remdesivir does not alter the potency of SF2523 and does not interfere with the binding of SF2523 to BRD4. To confirm this interaction, we performed NMR titration experiments using ^15^N-labeled BRD4 domains and showed that remdesivir is incapable of binding to either BD1, BD2, or the ET domain of BRD4, as no significant chemical shift perturbations were observed in ^1^H,^15^N heteronuclear single quantum coherence (HSQC) spectra of these domains (**Fig. 6**). However, the subsequent addition of SF2523 to the NMR samples of BD1 and BD2 led to large resonance changes, indicating robust binding, and no binding of SF2523 was detected to the ET domain of BRD4.

**Fig 6.**
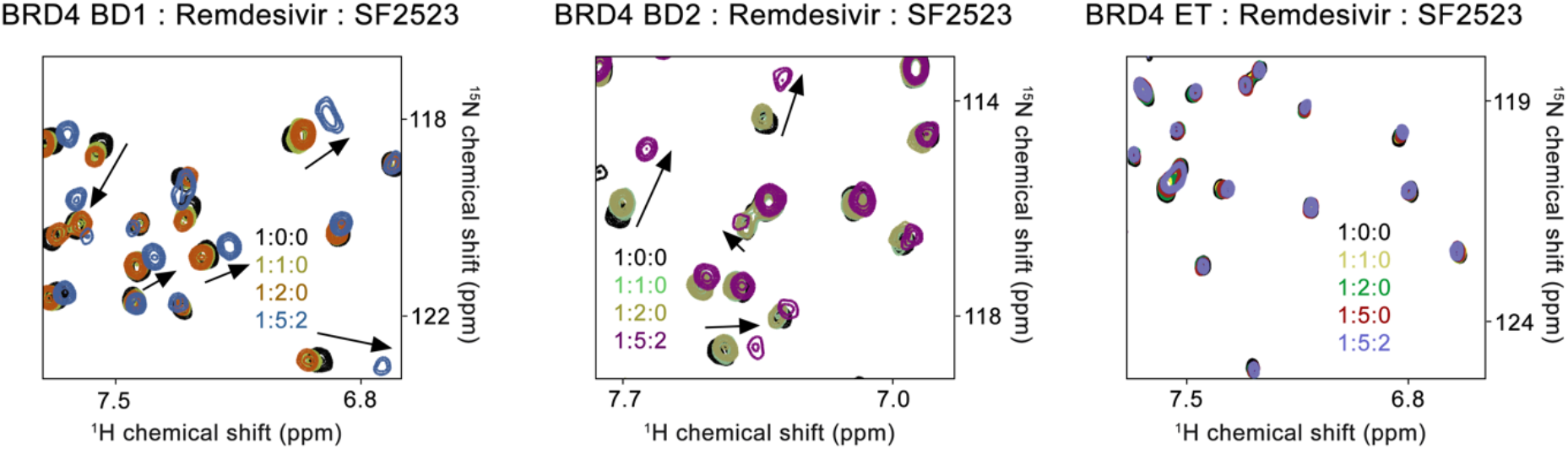
Remdesivir does not bind to the BRD4 BD domains and does not affect binding of SF2523 to BD_s_. Superimposed ^1^H,^15^N HSQC spectra of uniformly ^15^N-labeled BD1 (left), BD2 (middle) and ET (right) domains of BRD4 with the indicated ligands added in a stepwise manner. The spectra are color coded according to the protein:ligand ratios. Small resonance changes are due to weak binding of DMSO, which was confirmed by titrating DMSO alone to separate samples of these domains (data not shown).

## DISCUSSION

Other than remdesivir, a viral RNA-dependent RNA polymerase inhibitor (*34*), there are no confirmed efficacious molecularly targeted therapeutic agents for the treatment of COVID-19. Targeted therapies that exploit host-virus interaction are likely to be least impacted by the emergence of mutated strains of SARS-CoV-2 and produce more robust, durable treatment options. Gordon *et al*. identified two potential targets for SARS-CoV-2 infectivity, including BRD4 and mTOR kinase pathway (*29*). Using an *in vitro* we show that SF2523 is as potent and effective as remdesivir in blocking the SARS-CoV-2 cytotoxicity and infectivity at concentrations that are achievable and efficacious in animal models (*35*). SF2523 is highly selective and the only known dual inhibitor of PI3K-α/mTOR and BRD4 and is currently slated for clinical development. Hence, it is an excellent candidate for further development for systemic and pulmonary delivery for treatment of COVID-19. Importantly, SF2523 and multiple analogs are the product of extensive *in silico* molecular design to engineer chemotypes with dual or triple inhibitory properties against well-chosen synthetic lethality targets like BRD4, PI3K, and mTOR (*36-38*)

The ATP catalytic site of PI3K-α is driven by a critical hydrogen-bond interaction with the NH of Val851. The successful interaction of SF2523 as a PI3K inhibitor was designed with this mode of action where the oxygen atom of the morpholine group engages as a hydrogen-bond acceptor with the required NH of Val851. For the docking against PI3K, we initially created a p110-PI3K-α homology model using the crystal structure of human-derived PI3K-γ (PDB ID: 1E7V) where the human PI3K-α protein sequence was superimposed to the PI3K-γ protein sequence. The resulting 3D model of PI3K-α was subsequently used for docking of SF2523 where amino acids within a 10 Å radius at the ATP binding site were selected. Successful docking of SF2523 was achieved using FlexX, where the optimal bin36ding mode of SF2523 fulfills 2 pharmacophoric requirements, namely, the spatial location of the carbonyl group of SF2523 and a hydrogen-bond interaction between the morpholine group of SF2523 and Val851 (*39*). BRD4 bromodomain 1 (BRD4-BD1) active site recognizes acetylated lysine residues in histones via hydrogen-bond interaction with the NH of the conserved Asn140 residue. This binding site is a hydrophobic cavity that contains structurally conserved water molecules. For the docking of SF2523 against BDR4-BD1, a human-derived BDR4-BD1 crystal structure co-crystallized with small-molecule BRD4 inhibitor JQ1 (PDB ID: 3MXF) was obtained and amino acids within a 6 Å radius of JQ1 were selected. Using AutoDock Vina (*40*) and rDock (*41*), SF2523 was predicted to bind, forming a hydrogen-bond interaction between the carbonyl group of SF2523 and the NH of Asn140 and a conserved water molecule. The morpholine group of SF2523 is oriented towards the opening of the binding site (solvent-exposed). The subsequent successful crystallization of SF2523 with BRD4-BD1 (PDB ID: 5U28) confirmed the predicted binding interaction of SF2523 and it also revealed that one of the oxygen atoms of the benzodioxane moiety of SF2523 engages in 2 additional hydrogen-bond interactions with BRD4-BD1, one with GLN85 and one with another conserved water molecule (*35*). NMR data from our laboratory confirms interaction between BRD4 BD1 and acetylated lysine with the C terminus of SARS-CoV-2 E protein. Also, as per Zhang et al., other viral proteins have been shown to have functional interaction with BET BRD4 for viral pathogenesis (*42*).

From previous experience, it was observed that in patients infected with SARS and MERS, the viral load peaks only after a week of initial reporting of disease symptoms (*43, 44*). Whereas in the case of SARS-CoV-2, similar to influenza, the viral load peaks along with the onset of disease symptoms (*45*). It was reported that for influenza patients with high viral load at the beginning of disease symptoms, combinational antiviral therapy is more effective than monotherapies (*46, 47*). This is mirrored during the initial phase of the COVID-19 pandemic, where multiple combinational therapeutic approaches using several repurposed drugs show mild to moderate efficacy in treating severely ill patients (*48-50*). To extend this combinational therapeutic approach to treat COVID-19 patients *in vitro*, we tested combination treatment of SF2523 and remdesivir as described above. Our findings suggest that this drug combination is highly synergistic in the human bronchial epithelial cell line with 25-fold dose reduction for SF2523 and 4-fold dose reduction for remdesivir while maintaining similar therapeutic efficacy of individual monotherapies. This reduction in remdesivir dose is also expected to reduce the high incidence of adverse effects related to remdesivir monotherapy (*26, 51*).

To characterize the molecular basis for SF2523 mediated inhibition of SARS-CoV-2 replication and confirm the synergistic combination of this compound with remdesivir *in vitro*, we used NMR to look at the capacity of these two chemotypes to interact with BRD4-BD1, BRD4-BD2, and BRD4-ET domains. Consistent with the substantial synergy noted with the two agents, remdesivir did not bind to BRD4; however, SF2523 binds to both BD1 and BD2 but not the extra-terminal (ET) domain of BRD4, supporting differential molecular pathways targeted by these chemically distinct chemotypes. BRD2/BRD4 are epigenetic factors whose two N-terminal bromodomains (BDs) bind with acetylated histones and regulate several proteins (*52*). Several observations related to Gordon et al. (*29*) reporting that BRD4 binds to the SARS-CoV-2 E protein include: 1) the E protein forms a homopentameric cation-selective channel which is required for viral budding and release from cells, 2) the E protein is a 72-residue viroporin molecule, 3) mutations or deletions within the transmembrane channel attenuate viral pathogenicity (*53*). Data from emerging mutant strains of SARS-CoV-2 strains indicates presence of several variation in the C terminal domain E protein, that may impact its binding with host protein PALS1with altered disease pathogenesis (*54*). BRD2/BRD4 also has a significant role in cell cycle regulation, inflammation, and inhibition of BRD4 by SF2523 enhances the innate and adaptive immune response (*37*). Again, BRD4 drives the secretion of IL-6, CXCL8, and IL17A, which leads to the development of idiopathic pulmonary fibrosis and chronic obstructive pulmonary disease (COPD) (*55*). This suggests that BET inhibitors may disrupt the interaction between BRDs and E protein and will have a favorable outcome in treating COVID-19 patients. LARP1, a significant effector of the mTOR pathway, interacts with nucleocapsid protein (N) (*56*). LARP1 knockdown of inhibitors of mTOR inhibits MERS replication and have an immunosuppressive function (*57*). These data suggest potential important interactions between the SARS-CoV-2 virion and the host machinery elements essential for viral pathogenesis. Campbell et al., reported that SF2523 locks HIV replication in human cells via induction of autophagy (*58*). Moreover, Joshi et al. reported that SF2523 has a potential immunostimulatory impact on innate and adaptive T-cell immunity (*37*). Because SF2523 effectively targets two crucial elements of SARS-CoV-2 driven pathogenesis, BETs BRD2/BRD4, and mTOR, and because of its immune activation activity, SF2523 can be a potential therapeutic agent to control COVID-19. It can provide significant clinical benefit in preventing profound COVID-19 disease evolution.

Our initial efforts to screen the efficacy of SF2523 monotherapy and combinational therapy against SARS-CoV-2 in the 2D-cellular model is an essential step for initial screening of drug like compounds before initiation of pre-clinical studies and commonly used to explore the molecular mechanism of action. Our future studies will reveal their efficacy in pre-clinical animal models of COVID-19. Its synergistic effect will result in a favorable shift in the plasma Cmax/EC_90_ ratio. Ongoing structural studies to elucidate the interaction of SF2523 with host factors and viral targets will clarify mechanisms of efficacy, and potential resistance as the virus continues to undergo mutagenesis and selection during the pandemic. Considering the emerging data on the life cycle and biology of SARS-CoV-2 infection in humans and the rapid emergence of mutant variants resistant to vaccine candidates, we suggest that a combinatorial approach based on one small molecule which inhibits the interaction of multiple viral targets with host factors necessary for viral pathogenesis will be required to control the mortality and morbidity associated with the pandemic and the likely endemic phase of this disease globally.

In conclusion, our study demonstrates the *in vitro* efficacy of SF2523 as a monotherapy and in combination with remdesivir against SARS-CoV-2 infection. Considering the marked synergistic effect of these two molecules in a wide dose range in *in vitro* assays, we conclude that, SF2523 alone or in combination with remdesivir represents a future therapeutic approach to obviate the severe disease associated with the ongoing pandemic of SARS-CoV-2.

## MATERIALS AND METHODS

### Study design

In this *in vitro* study using a human bronchial epithelial cell line (UNCN1T) and Vero-STAT1 knockout cells, we evaluate the efficacy of SF2523 (PI3K-mTOR/BRD4 inhibitor) as a monotherapy or in combination with remdesivir against SARS-CoV-2. Using NMR, we also screen the interaction between different domains of BRD4 (BD1, BD2, and ET) and SF2523/remdesivir.

### Reagents, cell lines

Remdesivir (GS-5734) was obtained from Selleck Chemicals LLC, 14408 W Sylvanfield Drive, Houston, TX 77014 USA. Vero E6 (ATCC® CRL-1586™) and Vero-STAT1 knockout cells (ATCC® CCL-81-VHG™) were cultured in DMEM containing 10% fetal bovine serum, 2 mM L-glutamine, penicillin (100 units/ml), streptomycin (100 units/mL), and 10 mM HEPES. UNCN1T cells (a human bronchial epithelial cell line; Kerafast; cat# ENC011) were cultured using BEGM media (Bronchial Epithelial Cell Growth Medium; Lonza: cat# CC-3170) in FNC (Athena Enzyme Systems; cat# 0407) coated plates. All other molecular biology grade fine chemicals used in the study were purchased from Sigma-Aldrich, St. Louis, MO, USA, unless otherwise mentioned.

### *In silico* docking of SF2523 against PI3K-α and BRD4-BD1

The docking of SF-2523 with PI3K-α is described previously (*39*). *In silico* docking of SF2523 against BRD4-BD1 was performed using AutoDock Vina (*40*) and rDock (*41*). Crystal structure of BRD4 (PDB ID 3MXF) was downloaded from the Protein Data Bank (*59*), residues and water molecules within 6 Å^3^ of JQ1 were kept, hydrogen atoms and charges were added, and SF2523 was docked using default parameters. A minimum of 150 docking solutions were recorded and the conformation of SF2523 matching the expected binding mode (NH of Asn140 forming a hydrogen bond with the carbonyl group of SF2523) with the strongest predicted binding energy was selected.

### Protein expression and purification

The pGEX6P-BRD4 BD1 (amino acids 43-180) and pGEX4T-1 BRD4 BD2 (amino acids 342-460) plasmids were expressed in Escherichia coli BL21 (DE3) RIL cells grown in M9 minimal media supplemented with ^15^NH_4_Cl and purified as previously described (*35*). The bacteria cells were harvested by centrifugation, resuspended in 10 mM HEPES pH 7.5, 150 mM NaCl, 1 mM TCEP supplemented with DNase I and lysed by sonication. The BRD4 BD1 and BD2 constructs were purified from the cleared lysate using glutathione Sepharose 4B beads and the GST tag was cleaved with PreScission or thrombin protease. The protein was purified using a S100 column (GE Healthcare) equilibrated in 10 mM HEPES pH 7.5, 150 mM NaCl, 1 mM TCEP. BD1 protein fractions were assessed for purity (95%) by SDS-PAGE, buffer exchanged into 1X PBS (37 mM NaCl, 2.7 mM KCl, 4.3 mM Na2HPO4, 1.47 mM KH_2_PO_4_ supplemented with 5 mM DTT and concentrated to ∼10-20 mg/mL using 3 kDa Millipore concentrator and stored at -80 °C.

The pET32a-BRD4 ET domain (amino acids 600-700) vector was obtained from Timothy McKinsey. The N-terminal TRX-6His-S-tagged ET domain was expressed in Escherichia coli Tuner (DE3) competent cells using M9 minimal media supplemented with ^15^NH_4_Cl. Cells were induced with 0.5 mM IPTG, cultured for 16 h post-induction at 18 °C, and harvested at 5000 rpm. The His-tagged protein was purified with Nickel-NTA resin (Thermo Scientific) in 20 mM Sodium Phosphate (pH 7.0) buffer, supplemented with 500 mM NaCl, 10 mM imidazole and 2 mM BME. The tagged protein was eluted using a 30-300 mM imidazole gradient elution. The eluted protein was buffer exchanged into cleavage buffer containing 20 mM Tris pH 8.0, 50 Mm NaCl, 2 mM CaCl2 and the TRX-6His-S-tag was cleaved overnight at RT with enterokinase protease (NEB). The ET domain was further purified using Nickel-NTA resin (Thermo Scientific) to remove the cleaved tag. Size-exclusion chromatography was performed using a Superdex S75 column. The purified protein was buffer exchanged into 20 mM Tris pH 7.0, 100 mM NaCl, 5 mM DTT and concentrated to ∼10 mg/mL using 3 kDa Millipore concentrator and stored at -80 °C.

### NMR Experiments

For BRD4 BD1 or BD2 experiments, samples containing 0.2 mM ^15^N-labeled BRD4 BD1/BD2 in 1X PBS, 5 mM DTT, and 10% D_2_O were prepared. For BRD4 ET experiments, samples containing 0.15 mM ^15^N-labeled BRD4 ET domain in 20 mM Tris pH 7.0, 100 mM NaCl, 5 mM DTT, and 10% D_2_O were prepared. Remdesivir (Selleck Chemicals) and SF2523 (SignalRx Pharmaceuticals, Inc.) were dissolved in 100% DMSO. Remdesivir was titrated into each NMR sample up to 5 molar equivalents. Subsequently, SF2523 was added at a 1:2 equivalence. Two-dimensional ^1^H,^15^N HSQC spectra were acquired on a 600 MHz Varian spectrometer fitted with a cryogenic probe at 298 K. Spectra were processed with NMRPipe.

### Production of SARS-CoV-2 stocks

SARS-CoV-2 isolate USA-WI1/2020 (BEI; cat# NR-52384) was procured from BEI and passaged in Vero-STAT1 knockout cells. The viral titer was determined using plaque assay (*60*). In brief, Vero E6 cells were seeded in 6-well plates. After 24 hrs, cells were washed with sterile 1X PBS. The viral stock was serially diluted and added to cells in duplicate with fresh media and the plates were incubated at 37 °C for 1 hour with occasional shaking every 15 minutes. Then 2 mL of 0.5% agarose in minimal essential media (MEM) containing 5% FBS and antibiotics was added per well. Plates were incubated at 37 °C for 72 hours. Then the cells were fixed with 4% paraformaldehyde overnight, followed by the removal of the overlay and then stained with 0.2% crystal violet to visualize plaque forming units (PFU). All assays involving live SARS-CoV-2 were performed in University of Nebraska Medical Center (UNMC) BSL-3 core laboratory setting and approved by the University of Nebraska Medical Center institutional Biosafety Committee (IBC) with the approval number 20-05-029-BL3.

### Measuring the antiviral potential of SF2523 and remdesivir against SARS-CoV-2

UNCN1T and Vero-STAT1 knockout cells were seeded in 96-well plates 24 hours before infection at 20,000 cells/well. Different concentrations of SF2523 and remdesivir (100 μM, 10 μM, 5 μM, 1 μM, 0.1 μM, 0.01 μM and 0.001 μM) were added to the cells 2 hours before infection. The cells were infected with 0.1 MOI of SARS-CoV-2 using Opti-MEM I reduced serum medium (Thermo Fisher, Cat#31985062) and incubated for 1 hour at 37 °C with 5% CO_2_. For positive control, cells were treated with same volume of DMSO equivalent to the volume of drugs added. Mock infected cells received only Opti-MEM I reduced serum medium. At the end of incubation virus inoculum was removed, cells were washed with 1X PBS 3 times and fresh media supplemented with same concentration of drugs was added. Culture supernatant was collected at 24 hrs and 48 hrs post infection. The SARS-CoV-2 viral load was quantified in culture supernatant using RT-QPCR with primer probes targeting E gene of SARS-CoV-2 using PrimeDirect Probe RT-qPCR Mix (TaKaRa Bio USA, Inc) and Applied Biosystems QuantStudio3 real-time PCR system (Applied Biosystems, Waltham, MA, USA) per the manufacturer’s instructions. Primers and probes used for SARS-CoV-2 RNA quantification were as follows: E_Sarbeco_F1: 5’ – ACAGGTACGTTAATAGTTAATAGCGT – 3’ (400 nM), E_Sarbeco_R2: 5’ – ATATTGCAGCAGTACGCACACA – 3’ (400 nM) and E_Sarbeco_P1: 5’ – FAM - ACACTAGCCATCCTTACTGCGCTTCG - BHQ1 – 3’ (200 nM) as recommended by WHO. The SARS-CoV-2 genome equivalent copies were calculated using quantitative PCR (qPCR) control RNA from heat-inactivated SARS-CoV-2, isolate USA-WA1/2020 (BEI; cat# NR-52347). The percentage inhibition of SARS-CoV-2 replication in SF2523 and remdesivir treated wells were calculated with respect to viral concentration in positive control wells that were treated with DMSO (considered 0% inhibition) and negative control wells (uninfected cells). IC_50_ values were calculated using four parameter variable slope sigmoidal dose-response models using Graph Pad Prism 8.0 software.

### Measuring the combinational antiviral potential of SF2523 and remdesivir

To determine possible combinational antiviral effect of SF2523 and remdesivir against SARS-CoV-2, we tested 5 (remdesivir) X 4 (SF2523) dose combinations in SARS-CoV-2 infected UNCN1T and Vero-STAT1 knockout cells. The percentage inhibition of viral replication for each dose combinations were determined by RT-QPCR as described above. In brief, the UNCN1T and Vero-STAT1 knockout cells were seeded in 96-well plates at a density of 20,000 cells/wells 24 hrs before infection. Two hours before infection the cells were treated with different combination doses of SF2523 and remdesivir, infected with 0.1 MOI of SARS-CoV-2. Twenty-four hours post infection culture supernatant was collected and SARS-CoV-2 viral load was quantified using RT-QPCR as described above. The percentage inhibition of SARS-CoV-2 replication in the culture supernatant by different combined doses of SF2523 and remdesivir were determined with respect to viral concentration in positive control wells that were treated with DMSO (considered 0% inhibition) and negative control wells (uninfected cells). Then the percent inhibition of viral replication for 1:1 fixed dose combination of the drugs was used to generate CI-Fa, isobologram and dose-response plots. The combination index (CI) was calculated using multiple drug effect equation developed by Chou and Talalay using the CompuSyn algorithm (https://www.combosyn.com). A CI values of <1 indicate synergy, values >1 indicate antagonism, and values equal to 1 indicate additive effects (*31, 33, 61*). Dose-response percent inhibition matrix of single and combined treatment of remdesivir and SF2523 in SARS-CoV-2 infected UNCN1T and Vero-STAT1 knockout cells 24 hrs post infection and 3-D interaction landscape between remdesivir and SF2523 were calculated based on Loewe additive model using SynergyFinder v.2 (*62*).

### Statistical analysis

IC_50_ values were calculated using four parameter variable slope sigmoidal dose-response models using Graph Pad Prism 8.0 software. The combination index (CI) was calculated using multiple drug effect equation developed by Chou and Talalay using the CompuSyn algorithm (https://www.combosyn.com). 3-D interaction landscape between remdesivir and SF2523 was calculated based on Loewe additive model using SynergyFinder v.2.

## Acknowledgements

We acknowledge the UNMC BSL-3 core facility for allowing us to perform all *in vitro* experiments involving SARS-CoV-2. ‘The University of Nebraska Medical Center BSL-3 Core Facility is administrated through the Office of the Vice Chancellor for Research and supported by the Nebraska Research Initiative (NRI).’ “The following reagent was deposited by the Centers for Disease Control and Prevention and obtained through BEI Resources, NIAID, NIH: (a) SARS-Related Coronavirus 2, Isolate USA-WI1/2020, NR-52384 and (b) Quantitative PCR (qPCR) Control RNA from Heat-Inactivated SARS-Related Coronavirus 2, Isolate USA-WA1/2020, NR 52347.”

## Funding

This work is partially supported by National Institute of Allergy and Infectious Diseases Grant R01 AI129745, 5P30 CA036727-33 and Frances E. Lageschulte and Evelyn B. Weese New Frontiers in Medical Research Fund to SNB and NIH grants HL151334 and CA252707 to T.G.K; and CA215651 to DLD.

## Author contributions

Conceptualization: DLD, SNB. Methodology: AA, SNB, TGK, SNB.

Investigation: AA, KP, MT, KRV

Data analysis: AA, KP, MT, KRV, KBC, TGK, DLD, SNB

Funding acquisition: SNB

Writing – original draft: AA, SNB, DLD, TGK, GAM

Writing – review & editing: AA, KP, MT, KBC, KRV, TGK, DLD, SNB, GAM.

## Competing interests

DLD and GMA declare financial interest in SF2523 via their holding in SignalRx Pharmaceuticals, Inc.

## Data and materials availability

All data related with this study are presented in the manuscript or the supplementary materials.

**Fig S1.**
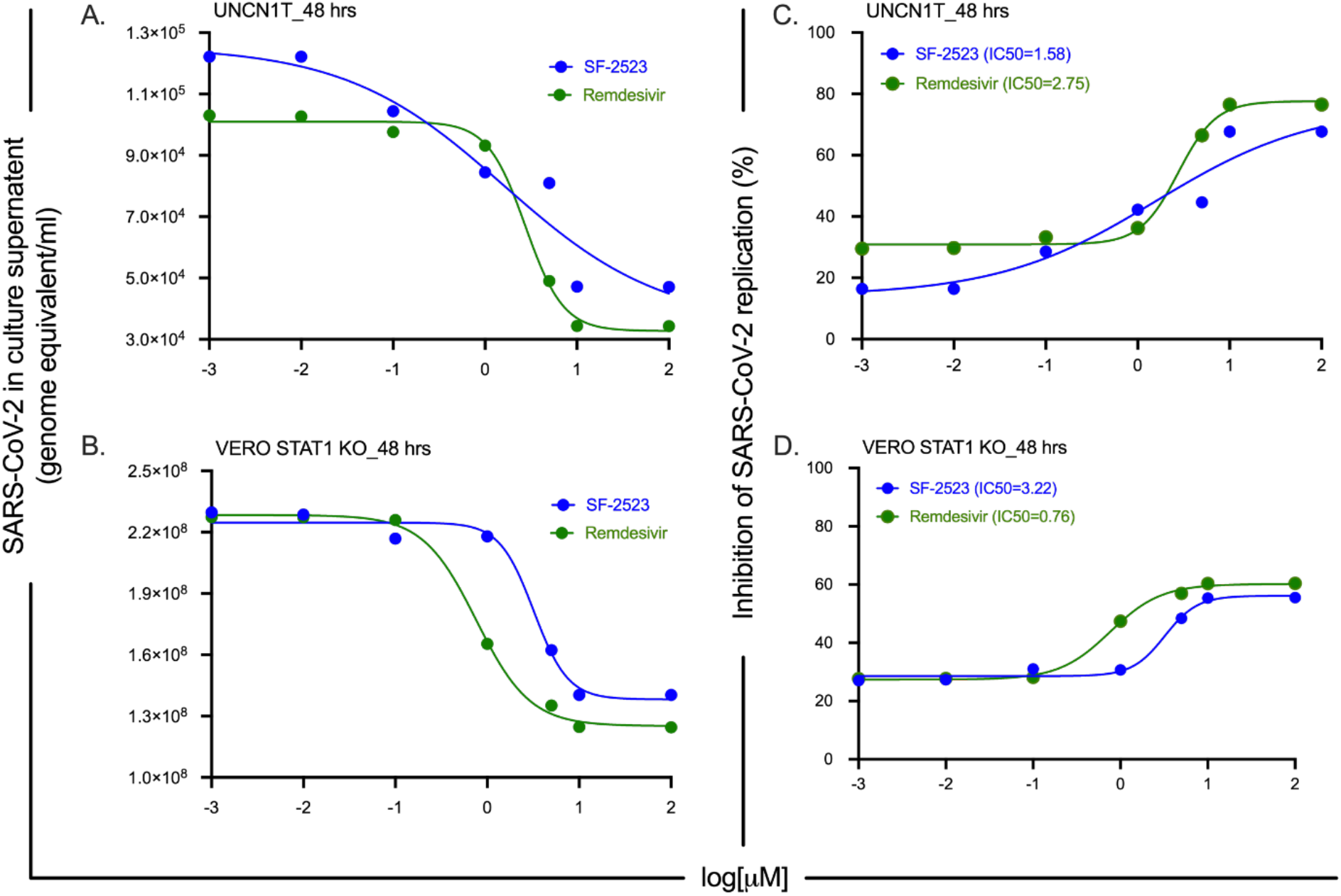
SARS-CoV-2 viral loads and dose-response curve in SF2523 and remdesivir treated and SARS-CoV-2 infected UNCN1T cells and Vero-STAT1 knockout cells. (**A & B**) Real-time quantitative PCR analysis of SARS-CoV-2 genome equivalent per ml of culture supernatant after 48 hrs post-infection in SF2523 (in blue) and remdesivir (in green) treated UNCN1T cells and Vero-STAT1 knockout cells with indicated drug concentrations respectively. (**C & D**) SF2523 (in blue) and remdesivir (in green) dose response curve by percentage inhibition of SARS-CoV-2 replication 48 hrs post infection in UNCN1T cells and Vero-STAT1 knockout cells with indicated drug concentrations respectively. In UNCN1T cells, SF2523 has an IC_50_ of 1.58 μM and remdesivir has an IC_50_ of 2.75 μM; In Vero-STAT1 knockout cells, SF2523 has an IC_50_ of 3.22 μM and remdesivir has an IC_50_ of 0.76 μM.

